# The impact of low input DNA on the reliability of DNA methylation as measured by the Illumina Infinium MethylationEPIC BeadChip

**DOI:** 10.1101/2021.12.22.473840

**Authors:** Sarah Holmes Watkins, Karen Ho, Christian Testa, Louise Falk, Patrice Soule, Linda V. Nguyen, Sophie FitzGibbon, Catherine Slack, Jarvis T. Chen, George Davey Smith, Immaculata De Vivo, Andrew J. Simpkin, Kate Tilling, Pamela D. Waterman, Nancy Krieger, Matthew Suderman, Caroline Relton

## Abstract

**Background:** DNA methylation (DNAm) is commonly assayed using the Illumina Infinium MethylationEPIC BeadChip, but there is currently little published evidence to define the lower limits of the amount of DNA that can be used whilst preserving data quality. Such evidence is valuable for analyses utilising precious or limited DNA sources.

**Materials and methods:** We use a single pooled sample of DNA in quadruplicate at three dilutions to define replicability and noise, and an independent population dataset of 328 individuals (from a community-based study including US-born non-Hispanic Black and white persons) to assess the impact of total DNA input on the quality of data generated using the Illumina Infinium MethylationEPIC BeadChip.

**Results:** Data are less reliable and more noisy as DNA input decreases to 40ng, with clear reductions in data quality; however samples with a total input as low as 40ng pass standard quality control tests, and we observe little evidence that low input DNA obscures the associations between DNAm and two phenotypes, age and smoking status.

**Conclusions:** DNA input as low as 40ng can be used with the Illumina Infinium MethylationEPIC BeadChip, provided quality checks and sensitivity analyses are undertaken.

## Introduction

Illumina Infinium MethylationEPIC BeadChips have been used extensively in epigenetic studies. Although Illumina recommend using at least 250ng of DNA on their BeadChips, there has been little published work examining the possibility of using less DNA than this. As DNA methylation (DNAm) profiling becomes more widespread, there is a need to ensure robust and reliable data can be generated from precious (e.g. clinical or historic) or limited (e.g. archaeological) biosamples. Two previous studies have assessed the effect of low levels of input DNA on the Illumina Infinium HumanMethylation450 BeadChip by generating data from multiple dilutions of the same biological samples. The first reported that correlations between genome-wide DNAm profiles remain above 0.96 for dilutions containing as little as 10ng of DNA (Whalley et al., 2021); the second reported correlations with input of 1μg for total input as low as 10ng remained above 0.92 (Hovestadt et al., 2013). However, no study has yet investigated the expected increase in signal variability or noise induced by low input DNA and its impact on statistical power to detect associations with DNAm; this is important because a number of studies have demonstrated that many probes on these BeadChips have low reliability, particularly where DNAm sites are either highly methylated or unmethylated and have low variance (Dugue et al., 2016; Forest et al., 2018; Xu & Taylor, 2021), and conceivably this might be exacerbated by low levels of input DNA applied to the BeadChip. Additionally, no comparison of data generated using different input levels has yet been carried out using a large population dataset.

Here we study assess whether low yields of input DNA are sufficient to reliably detect associations with DNA methylation measured using the Illumina Infinium MethylationEPIC BeadChip. The study consists of two parts: an initial analysis, where we assess reliability and noise within a single sample at three DNA concentrations; and a subsequent assessment of total input DNA on data quality and phenotype associations, using an independent population-based DNAm dataset of 328 individuals from the My Body My Story (MBMS) study (Krieger et al., 2011). We believe this is the first study assessing the impact of low input DNA explicitly utilising data from a large and socially diverse cohort.

## Materials and methods

### Study participants

The initial analysis (which we refer to as Study 1) included varied DNA dilutions from a single source, utilising a DNA sample pooled from several individuals stored at −80°C. Unfortunately no details about the individuals contributing to this pooled sample were available. The sample was used to generate three dilutions resulting in three quantities of total DNA input (40ng, 200ng, and 400ng), in quadruplicate, resulting in 12 samples.

The second analysis (which we refer to as Study 2) utilised the MBMS cohort. MBMS is a cohort recruited from four community health centers in Boston between 2008 and 2010, and was designed to investigate how racial discrimination affects risk of cardiovascular disease, taking into account a range of social and environmental factors. The cohort and recruitment procedures have previously been described in detail (Krieger et al., 2011); briefly, the study recruited 1005 individuals who met study inclusion criteria and were randomly selected from the patient rosters of the community health centers. Participants were eligible if they were aged between 35 and 64 years, had been born in the US, and self-identified their race/ethnicity as white non-Hispanic or black non-Hispanic. Among the 1005 MBMS participants, 85% provided a finger prick blood sample on to filter paper (409 black; 466 white), and consequently biological material was limited and in some instances of poor quality. Blood spots were stored at −20°C, and DNA was extracted from blood spots using the QIAamp DNA Investigator Kit for FTA and Guthrie cards, with samples randomised across 96 well plates. Of the 875 participants who provided blood spots, 472 of the samples were judged to be suitable for DNA extraction (blood spots judged not to be suitable were primarily the first community health center where recruitment took place, whose membership was predominantly white). Of those, 48 yielded less than 40ng of DNA, the lowest input level investigated in Study 1, so we removed them from further analysis. After removing a further 96 participants from the sample set due to poor quality DNA extraction (as determined by high numbers of undetected probes on the EPIC BeadChip), there were 328 participants with DNA methylation data for analysis. DNAm data were generated using the Illumina Infinium MethylationEPIC BeadChip as described below.

### DNA methylation data generation

For both studies, extracted DNA was bisulphite converted with the EZ DNA Methylation-Lightning™ Kit (Zymo Research) according to the manufacturer’s instructions. The eluant from the bisulphite-converted DNA was then applied to the Illumina Infinium MethylationEPIC Beadchip to measure DNA methylation, according to the manufacturer’s protocol. The EPIC BeadChips were scanned using Illumina iScan, with an initial quality review conducted with GenomeStudio. Sample QC and normalisation were conducted using the pipeline implemented in the *meffil* R package, which has previously been described in detail (Min, Hemani, Davey Smith, Relton, & Suderman, 2018). Blood cell composition was estimated for MBMS using a deconvolution algorithm (Houseman et al., 2012) implemented in *meffil*, based on the “blood gse35069 complete” cell type reference. DNA methylation is reported in beta values; this measures methylation on a scale of 0 (0% methylation) to 1 (100% methylation).

### Study 1: Assessing reliability of DNAm measurement with low input DNA

Using the single pooled sample of DNA described above, we used two methods to assess the reliability of DNAm measurements at different input DNA levels. Firstly, we assessed how well the measurements at the lower input levels (200ng and 40ng) replicate the measurements obtained with 400ng input DNA. To do this we calculated the mean methylation at each DNAm site across the four technical replicates at each input level. We then partitioned DNAm sites into bands based on their methylation level measured at 400ng (used as the reference level) in increments of 5%. Within each partition we calculated the standard deviation of the DNA methylation levels across all sites in the partition and visualised this variation using boxplots at 40ng and 200ng. Stronger replication of the 400ng measurements would correspond to smaller variation within each partition.

Secondly, we assessed the noise in DNAm measurement within each of the three DNA input levels using their four replicates. At each DNAm site, we took the mean of replicates 1 and 2, and used these means to partition the dataset into bands of 5% methylation as we did for the first analysis. Within each partition we then calculated the mean of replicates 3 and 4 at each DNAm site. We visualised the variation within each partition using boxplots of the mean of replicates 3 and 4 for all sites within the partition. Levene’s test (*leveneTest* in the R package car) was used to determine whether lower DNA input was associated with greater variance within each partition. Greater measurement noise would correspond to greater variance. As we tested 20 partitions, we used a p-value threshold corrected for multiple tests (p<0.05/20).

### Study 2: Assessing the impact of low input DNA in a cohort study

We then assessed how low DNA input affects the quality of Illumina Infinium MethylationEPIC Beadchip data using data from our cohort study, MBMS. We conducted two sets of analyses: we calculated a variety of QC-related metrics, and evaluated the effect of input DNA level on robust associations that have been reported in the DNAm literature.

We utilised two standard QC metrics to represent data quality: proportion of probes with low signal, and median methylated signal across all probes on the BeadChip. Low signal was assessed using detection p-values, which indicate confidence that the signal from a probe is detectable above background noise. We used a detection p-value threshold of 0.01 to distinguish between detection success and failure. We plotted the relationship between the number of undetected probes and DNA input level and correlated the two variables to test the strength of the association. Median methylated signal refers to the strength of probe signal due to binding of methylated DNA to a probe. We plotted median methylated signal per sample against DNA input level, and tested their association.

In addition to these QC steps, we compared DNAm measurements for each sample against a gold standard derived from all 135 samples with DNA input >200ng by simply calculating the mean for each individual probe on the BeadChip across the 135 samples. For all remaining samples with DNA <200ng (n=193), we calculated the difference between the methylation value at each probe and that of the gold standard, and summarised these differences per sample by taking the mean absolute difference, or MAD. We then evaluated the association between MAD and DNA input level using plots and by calculating Pearson’s correlation.

We tested whether variance in DNAm is associated with DNA input level at each site on the BeadChip using a procedure detailed elsewhere (Staley et al., 2021). We firstly use the function *rq* (a least absolute deviation regression) from the R package *quantreg* to test the association between methylation at each cpg site and DNA input level, including batch, cell counts, age, gender, smoking, and BMI as covariates in the model. From this model we take the absolute values of the residuals, and then test for an association between those residuals and DNA input level. We extracted coefficients and p-values from the model and applied a Bonferroni-corrected threshold of 5.8e-08 (0.05/857774) to identify associated sites. We took the −log10 of the p-values and created a Manhattan plot.

To assess how DNA input level might affect downstream analyses, we tested whether lower input DNA might increase noise in DNAm measurements to the point of obscuring associations with phenotypes. We tested this for two phenotypes, age and smoking status, because these both have robust associations with DNAm that can be reliably detected. We estimated epigenetic age using the Horvath (Horvath, 2013) and Hannum (Hannum et al., 2013) clocks. The absolute of the difference between epigenetic age and chronological age was calculated, and this difference was plotted against DNA input level. As random measurement error in a continuous outcome increases standard error, we used loess regression to test whether the standard error increased as DNA input decreased - this asks whether epigenetic age prediction becomes more noisy as DNA input level decreases. To assess whether noise might obscure the relationship between DNAm and smoking status, we tested whether reduced DNA input was associated with increased noise in the AHRR CpG cg05575921. To do this we regressed out the effect of smoking status on the unadjusted value of cg05575921 using a linear model, and tested the relationship between the absolute values of the residuals from the model and DNA input level in the same way as for epigenetic age, using loess regression. To derive smoking status, MBMS participants were asked “Have you smoked 100 or more cigarettes in your entire life?” and “Do you now smoke cigarettes every day, some days, or not at all?”; responses were combined and consolidated as either ‘current smoker’, ‘former smoker’ or ‘never smoker’.

## Results

### Participant characteristics

Three quarters (74%) of participants in our study identified their race/ethnicity as Black non-Hispanic, 56% lived in areas with high numbers of individuals below the poverty line, and two thirds (66%) had less than 4 years of college education. Characteristics of the 328 participants are summarised in Table 1; DNA quantity is marginally associated with smoking status (lower quantities for former and never smokers compared to current smokers), race/ethnicity (lower quantities for white participants), and education (lower quantities for participants with <4 years of college education, and highest quantities for participants with less than high school education).

**Table 1:** Characteristics of the 328 MBMS participants with DNAm data passing QC.

|  |  | **N (%) unless otherwise stated (total = 328)** | **Regression coefficient/ Mean input DNA (ng)** | **Association with total input DNA (p value)** |
| --- | --- | --- | --- | --- |
| **Age** | Mean (years) | 48.9 (mean) | 0.05 | 0.97 |
|  | Standard deviation (years) | 7.9 (SD) |  |  |
| **Gender** | Women (cis-gender) | 210 (64%) | 231.7 ng | reference |
|  | Men (cis-gender) | 118 (36%) | 201.1 ng | 0.13 |
| **Smoking** | Current | 150 (46%) | 248.7 ng | reference |
|  | Former | 63 (19%) | 189.9 ng | 0.03 |
|  | Never | 115 (35%) | 201 ng | 0.03 |
| **Race/ethnicity** | Black NH | 242 (74%) | 234.2 ng | reference |
|  | White NH | 86 (26%) | 182.7 ng | 0.02 |
| **Census tract poverty, % (2005-2009)** | <5% below poverty line | 17 (5%) | 230.2 ng | reference |
|  | >=5%,<10% below poverty line | 53 (16%) | 246.1 ng | 0.75 |
|  | >=10%,<20% below poverty line | 75 (23%) | 176.4 ng | 0.26 |
|  | >=20%,<40% below poverty line ("poverty area") | 131 (40%) | 231.7 ng | 0.97 |
|  | >=40% below poverty line ("extreme poverty area") | 52 (16%) | 227.7 ng | 0.96 |
| **Education** | Less than high school | 42 (13%) | 273.7 ng | 0.003 |
|  | > High school, < 4 years college | 218 (66%) | 226.2 ng | 0.02 |
|  | 4+ years college | 68 (21%) | 170.3 ng | reference |

### DNA methylation data

In Study 1, quality control identified 55,706 probes for removal due to failed detection, primarily for to samples with 40ng input DNA (see Figure 1). This left 807,787 CpG sites for further analysis. For MBMS (Study 2) a total of 328 samples and 857,774 sites passed probe detection checks; however 35 of these samples had a mismatch between the gender they reported in the study and sex as predicted by probe signal intensities targeting sites on the X and Y chromosomes. Furthermore, whereas correlation between chronological age and age estimated from DNA methylation was high in the dataset (Horvath clock R=0.63, Hannum clock R=0.69), correlation among these 35 was very low (Horvath clock R=-0.01, Hannum clock R=0.18). We included these 35 samples in our assessment of data quality using QC analyses as they displayed no evidence of low quality, and there was no relationship between sex/gender mismatch and DNA concentration (p=0.72, Wilcoxon rank sum test); but they were removed from analyses with phenotypes (epigenetic age and smoking), leaving 293 individuals in the phenotype analysis.

**Figure 1:**
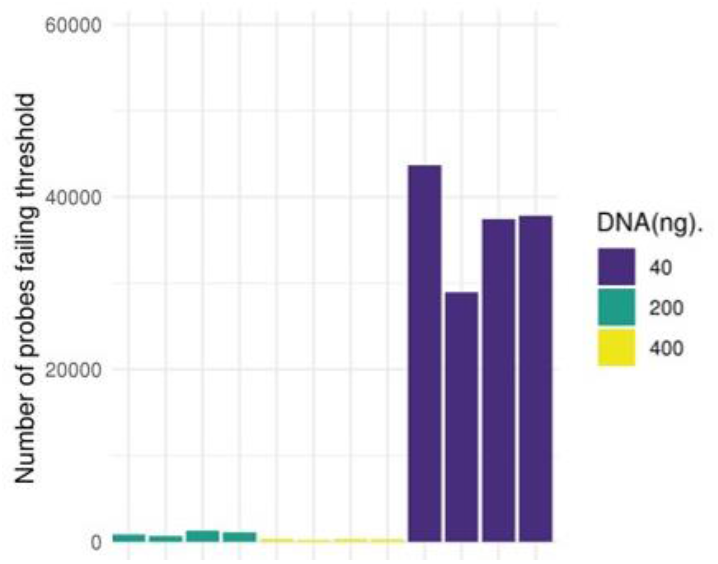
Number of probes that fail the detection p-value at 40ng, 200ng and 400ng. Each bar represents one sample.

### Study 1 results: Assessing reliability of DNAm measurement with low input DNA

The overall distributions of the methylation measurements across the BeadChip are virtually identical at 200ng and 400ng of input DNA, but it is skewed toward higher methylation levels for the 40ng dilution (Figure 2A). To investigate the reliability of methylation measurements when samples have low input DNA, we assessed how well measurements at 200ng and 40ng replicated those at 400ng by binning methylation sites according to methylation levels determined at 400ng, our reference. For both 40ng and 200ng, variance within each bin tends to be larger in bins representing intermediate methylation levels at 400ng. The main difference is the variation as measured by standard deviation tends to be 2-4 times larger at 40ng (SD = 0.01 to 0.04) than at 200ng (SD = 0.02 to 0.17) (Figure 2B and 2C; Supplementary Table 1). This indicates a reduced replication of 400ng signal at 40ng compared to 200ng.

**Figure 2:**
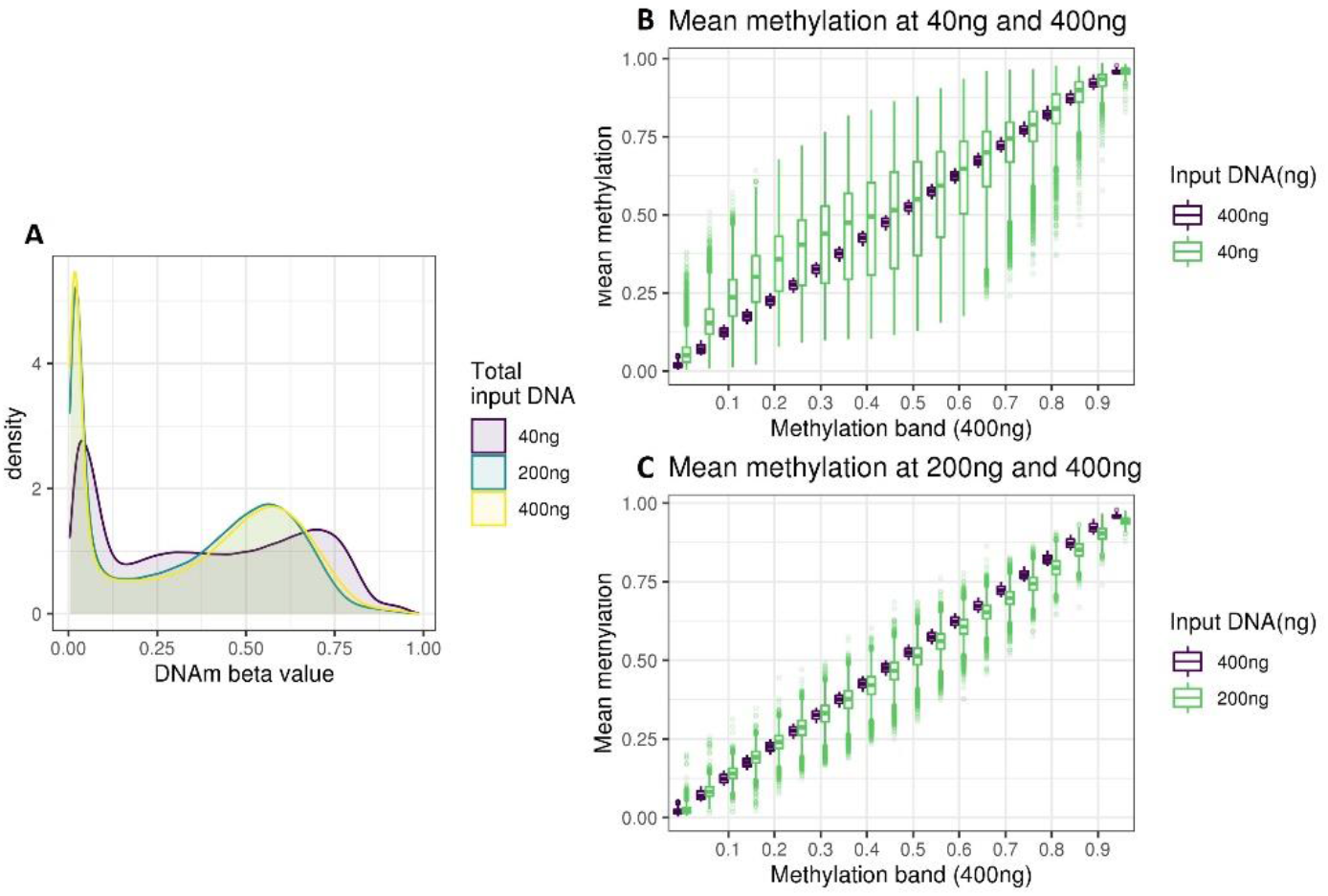
**A**: Density of methylation beta values across the EPIC BeadChip for 40ng, 200ng and 400ng DNA. **B**: boxplot of the methylation of DNAm sites at 40ng, grouped in bins of 0.05 based on the methylation level of the DNAm site at 400ng. **C**: boxplot of the methylation of DNAm sites at 200ng, grouped in bins of 0.05 based on the methylation level of the DNAm site at 400ng.

As we had quadruplicate measurements for the three DNA input levels we were also able to assess the noise within each input level. This is important because measurements by Illumina Infinium MethylationEPIC Beadchips are known to be noisy, and low concentrations of DNA may exacerbate this issue (Belsky et al., 2020; Xu & Taylor, 2021). To assess noise at each DNA input level, we used two replicates to partition methylation sites by methylation level and then calculated the variance of each bin from the other two replicates. Using Levene’s test of variance to compare bin variances between dilutions, we show that 200ng is noisier than 400ng in 17 out of the 20 partitions (at p<0.05/20); and that 40ng is noisier than both 400ng and 200ng in all 20 partitions (at p<0.05/20). This demonstrates that as DNA input level decreases, measurement noise increases. Bin variances are illustrated in Figure 3, and Levene’s test statistics in Supplementary table 2.

**Figure 3:**
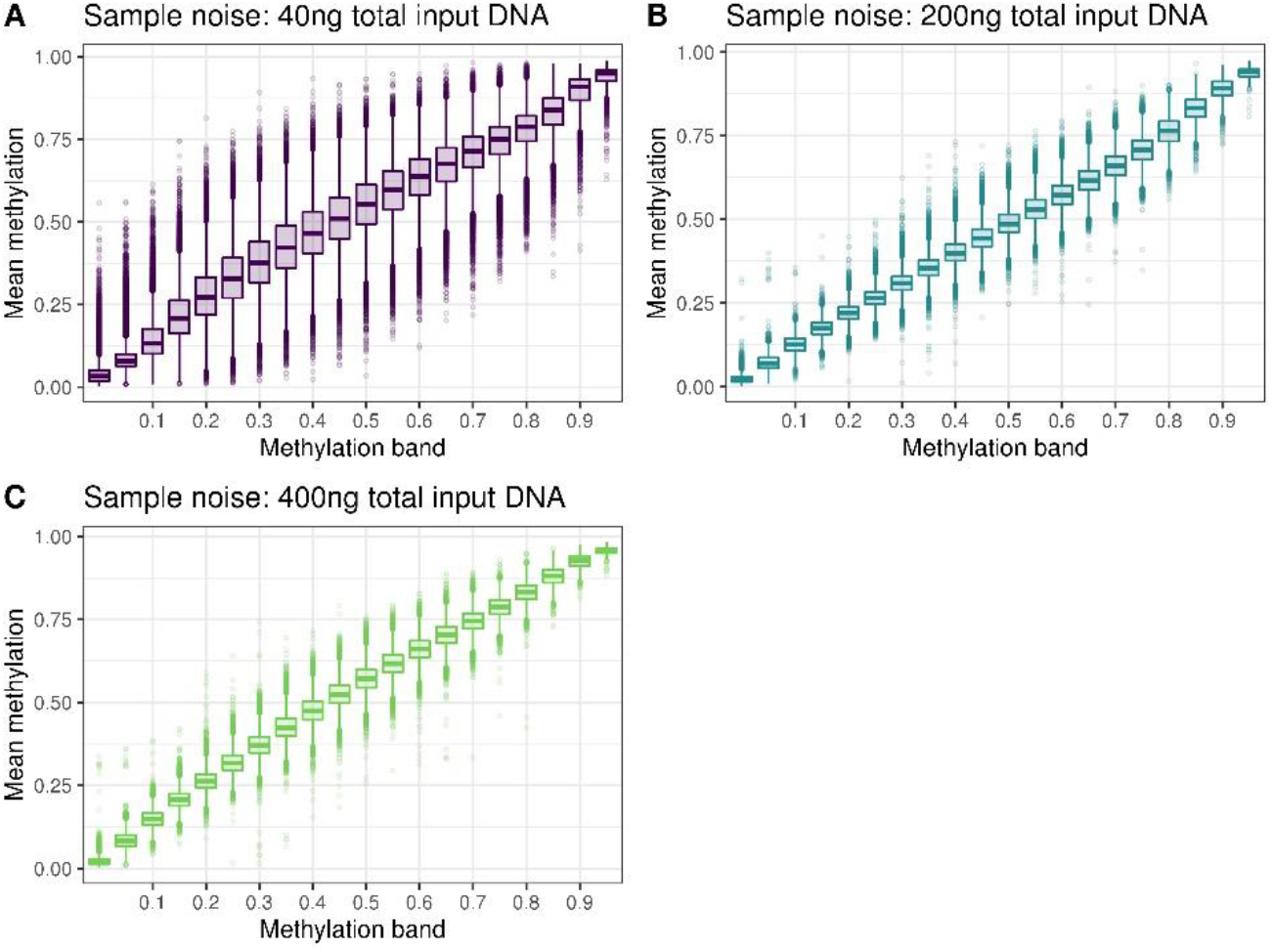
Plots of sample noise at **A** 40ng, **B** 200ng and **C** 400ng total input DNA. All CpG sites were binned into 5% partitions of methylation beta value based on the mean of replicates 1 and 2, and the mean of replicates 3 and 4 was used to create the boxplots.

### Study 2 results: Assessing the impact of low input DNA in a cohort study

DNA for the MBMS cohort was extracted from dried blood spots and resulted in a range of DNA quantities for the 472 participants for whom DNA was extracted (mean 173ng, range 0ng to 1186.8ng). As we excluded participants with less than 40ng DNA, and those with poor quality DNA, DNA quantities were higher for the 328 participants who were included in the analyses included in this paper (mean 220.7ng, range 40.6ng to 1186.8ng). We assessed the impact of DNA input level on the quality of the MBMS DNAm data in three ways: the proportion of probes failing detection p-value, the median strength of methylated signal, and the mean absolute deviation of samples with lower than recommended DNA (200ng) in comparison to a gold standard based on measurements from the samples with at least 200ng DNA. Although the proportion of undetected probes increases as input DNA decreases (R=-0.26, p=1.2e-06), we find that samples down to 40ng have acceptable proportions of undetected probes, i.e. within the limits of typical Illumina QC pipelines (<0.025%) (Figure 4A). Similarly, although median methylated signal is correlated with DNA input level (R=0.46, p<2.2e-16), the signal does not fall below typical quality thresholds (Figure 4B). Finally, as expected, samples with lower DNA input level tend to have higher mean absolute deviation from the gold standard based on samples with at least 200ng of DNA (R=-0.38, p=6.8e-08); Figure 4C. Thus, we have shown that measurement quality and precision decrease with input DNA levels.

**Figure 4:**
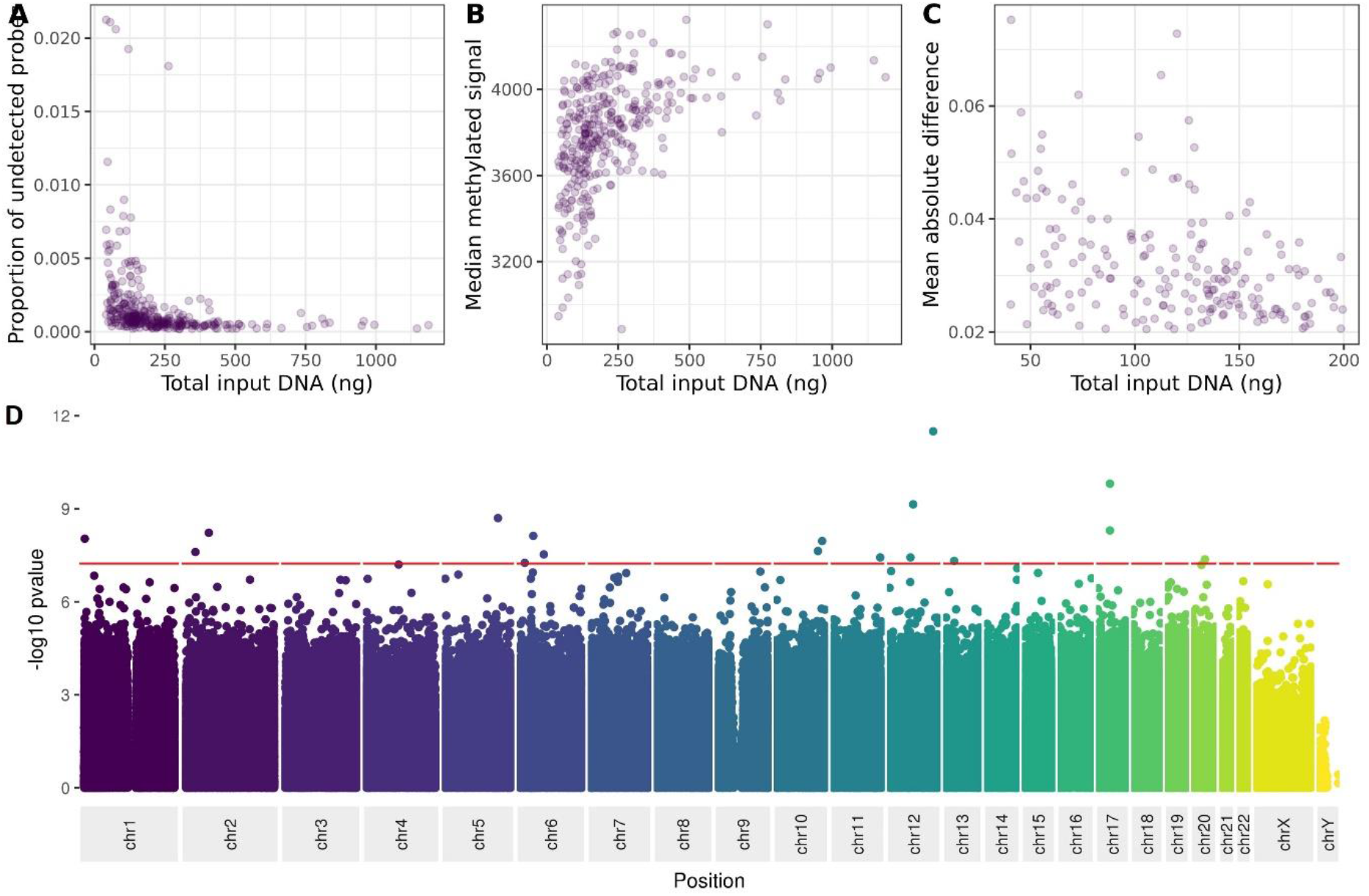
The relationship between DNA input level and **A** proportion of probes failing detection p-value, **B** median methylated signal, **C** mean absolute deviation from a composite of the high-input samples, **D** variance at each site on the Illumina Infinium MethylationEPIC Beadchip.

We then asked whether low DNA input level affects the variance of methylation measurements at specific individual sites on the BeadChip. Using linear regression with a BeadChip-wide Bonferroni-corrected threshold of 5.8e-08, we observe associations between variance in methylation value and DNA input level at 17 sites (Figure 4D and Supplementary table 3).

Finally, we asked to what extent DNA input level might affect our ability to detect the association between DNA methylation and two phenotypes (age and smoking status) that have been strongly associated with DNA methylation in numerous previous studies. Using loess regression we found no evidence of an increase of standard error with decreasing DNA input level, suggesting that low DNA input does not obscure associations between DNA methylation, and age estimation (as measured by the Horvath and Hannum epigenetic clocks) or smoking status (Figure 5).

**Figure 5:**
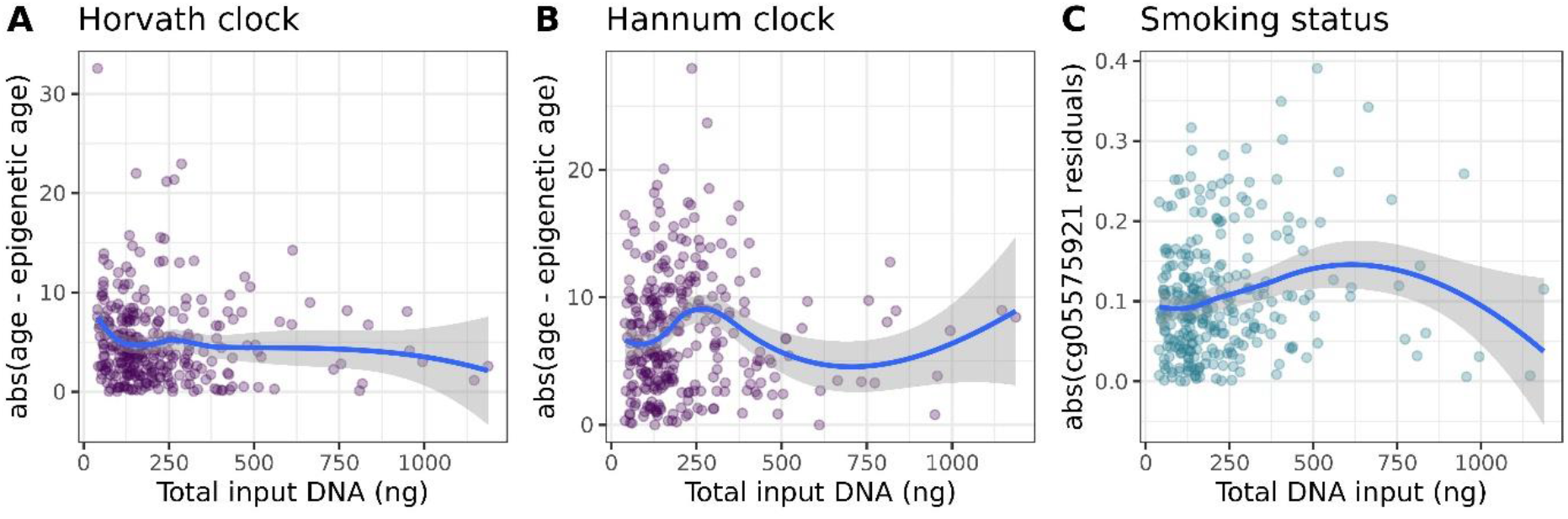
**A** scatter plot of the relationship between DNA input level, and the absolute of the difference between epigenetic age as estimated by the Horvath clock and chronological age; **B** A scatter plot of the relationship between DNA input, and the absolute of the difference between epigenetic age as estimated by the Hannum clock and chronological age; **C** A scatter plot of the residuals of the model lm(cg05575921 ~ smoking status) against total DNA input. For each plot the blue line represents the loess regression line and the grey shading represents the standard error of the regression.

## Discussion and conclusions

This study demonstrates that although as little as 40ng is sufficient to produce Illumina Infinium MethylationEPIC Beadchip DNAm data that passes standard QC checks, data quality and reliability diminish as DNA input decreases. However, this reduction in data quality may have limited practical impact on downstream analyses, as we show that two strong phenotype associations with DNAm – age and smoking – do not appear to be adversely affected by using DNA input levels as low as 40ng. We hope this demonstration can empower studies to conduct DNAm investigations where it might have previously been assumed that samples were too limited to provide sufficient DNA; but due to the increase in both noise and variance that we have demonstrated, we would recommend caution and use of sensitivity analyses when working with less than 200ng DNA on the Illumina Infinium MethylationEPIC Beadchip.

Our evaluation of DNA from a single source at three dilutions illustrates that using 40ng of DNA produces noisier measurements than using 200ng, and using 200ng is noisier than 400ng. This corresponds to reduced agreement we report between measurements at 40ng than at 200ng compared to those at 400ng. Analysis of data from a cohort of 328 individuals shows a clear impact of decreasing DNA input on the proportion of probes failing detection and on the strength of methylation signal; this is presumably because there is less DNA binding to probes. This also appears to be the reason for the clear impact of decreasing DNA input level on increasing deviation from a gold standard composite profile based on samples with at least 200ng DNA. Importantly, our analyses show how fast data quality decreases as input DNA decreases, so our findings can be used identify thresholds on input DNA suited to specific research questions. It is notable that data quality is acceptable as assessed by common quality control metrics when input DNA as low as 40 ng.

We would strongly recommend that researchers using DNA input of less than 200ng should run quality checks and sensitivity analyses with the lower concentration samples. As we show DNA input is strongly associated with variance at many specific DNAm sites, we would suggest extra caution around these sites as they may be particularly affected by low DNA concentrations. We have provided the full summary statistics from this variance EWAS in Supplementary table 3 so that researchers can utilise these results with p-value or effect thresholds appropriate to their data and research question. Finally, we show that associations of DNA methylation with age and smoking are observable in samples with input DNA as low as 40ng, and that we see little evidence of increased measurement noise as DNA input decreases to 40ng. This is encouraging as it suggests that useful data can be derived from low input DNA; but we would caution that we have tested two exposures that have particularly strong associations with DNAm, and so associations for other phenotypes may be affected to a greater extent.

Strengths of our study include complementary analyses of both control and human cohort DNA samples; and both the large number and social diversity of individuals in the cohort analysis (n=328). Social diversity is very important for study generalisability because DNAm is affected by our social environment. The main limitation is that the impact of DNA input level may well be different for differing sample types and provenances, as DNA quality is affected by storage and extraction methods. Indeed we can see that with the much larger number of probes failing detection p-value thresholds from the 40ng samples of pooled frozen DNA in comparison to samples with close to 40ng that were extracted from dried blood spots as part of the MBMS study. However, we were not able to measure DNA quality in this study so cannot comment further on how this may impact results. Additionally, we did not assay less than 40ng DNA, so we cannot comment on how data quality might be affected by lower levels of DNA input; future studies may want to investigate data quality using lower inputs.

## Acknowledgements

We are extremely grateful to all the MBMS participants who took part in this study, the staff at the four participating community health centers, and the researchers who conducted participant recruitment and interviews. We thank Elmer Freeman, Director of the Center for Community Health Education Research and Services, Inc (Northeastern University, Boston) and Brent Coull, Professor of Biostatistics and Environmental Health (Harvard TH Chan School of Public Health) for their involvement with the original MBMS study and continued engagement with this project.

